# The five *Urochloa* spp. used in development of tropical forage cultivars originate from defined subpopulations with differentiated gene pools

**DOI:** 10.1101/2021.07.21.453213

**Authors:** J Higgins, P Tomaszewska, TK Pellny, V Castiblanco, J Arango, J Tohme, T Schwarzacher, RA Mitchell, JS Heslop-Harrison, JJ De Vega

## Abstract

**Background and Aims:** *Urochloa* (syn. *Brachiaria*, and including some *Panicum* and *Megathyrus*) is a genus of tropical and subtropical grasses widely sown as forage to feed ruminants in the tropics. A better understanding of the diversity among *Urochloa* spp. allow us to leverage its varying ploidy levels and genome composition to accelerate its improvement, following the example from other crop genera.

**Methods:** We explored the genetic make-up and population structure in 111 accessions, which comprise the five *Urochloa* species used for the development of commercial cultivars. These accessions are conserved from wild materials from collection sites at their centre of origin in Africa. We used RNA-seq, averaging 40M reads per accession, to generate 1,167,542 stringently selected SNP markers that tentatively encompassed the complete *Urochloa* gene pool used in breeding.

**Key Results:** We identified ten subpopulations, which had no relation with geographical origin and represented ten independent gene pools, and two groups of admixed accessions. Our results support a division in *U. decumbens* by ploidy, with a diploid subpopulation closely related to *U. ruziziensis*, and a tetraploid subpopulation closely related to *U. brizantha*. We observed highly differentiated gene pools in *U. brizantha*, which were not related with origin or ploidy. Particularly, one *U. brizantha* subpopulation clustered distant from the other *U. brizantha* and *U. decumbens* subpopulations, so likely containing unexplored alleles. We also identified a well-supported subpopulation containing both polyploid *U. decumbens* and *U. brizantha* accessions; this was the only group containing more than one species and tentatively constitutes an independent “mixed” gene pool for both species. We observed two gene pools in *U. humidicola*. One subpopulation, “humidicola-2”, was much less common but likely includes the only known sexual accession in the species.

**Conclusions:** Our results offered a definitive picture of the available diversity in *Urochloa* to inform breeding and resolve questions raised by previous studies. It also allowed us identifying prospective founders to enrich the breeding gene pool and to develop genotyping and genotype-phenotype association mapping experiments.

**HIGHLIGHT:** We clarified the genetic make-up and population structure of 111 *Urochloa* spp. forage grasses to inform cultivar development.

## INTRODUCTION

*Urochloa* (syn. *Brachiaria*, and including some *Panicum* and *Megathyrus*) is a genus of tropical and subtropical grasses widely sown as forage to feed ruminants in the American and African tropics, particularly in areas with marginal soils. *Urochloa* grasses exhibit good resilience and low nutritional needs (Miles, 2007, Gracindo et al., 2014, Maass et al., 2015). Five species, *U. ruziziensis, U. decumbens, U. brizantha, U. humidicola*, and *U. maxima* are broadly used as fodder plants, covering over 100M hectares in Brazil alone. Such an enormous area, about half that of each of the most widely grown cereals, wheat or maize, has a huge environmental impact in terms of displacement of native species, water usage, and provision of ecosystem services. In addition to extensive pasture systems in Latin America, *Urochloa* is also planted in intensive smallholder systems in Africa and Asia (Keller-Grein et al., 1996, Maass et al., 2015). Breeding programmes in different countries have exploited the diversity among *Urochloa* spp. for the development of commercial forage cultivars by recurrent selection over many years (Jank et al., 2014, Tsuruta et al., 2015, Worthington and Miles, 2015).

The genus *Urochloa* includes species previously classified under *Brachiaria*, *Megathyrsus*, *Eriochloa* and *Panicum* (Torres González and Morton, 2005, Kellogg, 2015). Joint missions between 1984 and 1985 conducted by the CGIAR (Consultative Group on International Agricultural Research) centres in several African countries collected wild materials from the species in the genus, mostly as live plant cuttings or ramets (Keller-Grein et al., 1996). These activities built a global grass collection with ~ 700 *Urochloa* accessions that are held at CIAT (Centro Internacional de Agricultura Tropical), ILRI (International Livestock Research Institute), and EMBRAPA (Brazilian Agricultural Research Corporation).

Three *Urochloa* species (*U. brizantha*, *U. decumbens, and U. ruziziensis*) have been arranged in an agamic (apomictic) group or complex (Do Valle and Savidan, 1996, Renvoize et al., 1996, Ferreira et al., 2016, Triviño et al., 2017). Crosses between ~ 10 founders from these three species were completed in the late 1980s and their progeny constitutes the basis of the current recurrent selection breeding programmes at CIAT and EMBRAPA (Miles et al., 2006). On the other hand, *U. humidicola* and *U. dictyoneura* have been arranged in the “humidicola complex” (Lutts et al., 1991, Renvoize et al., 1996, Triviño et al., 2017). More recently, independent *U. humidicola* breeding programs have also been established at CIAT and EMBRAPA after the discovery in the mid-2000s of a natural sexual polyploid germplasm accession that could be crossed with other apomictic polyploid *U. humidicola* pollen donors (Jungmann et al., 2010, Vigna et al., 2011a).

*Urochloa* spp. show varying ploidy levels and sub-genome compositions (Do Valle and Savidan, 1996, Keller-Grein et al., 1996, Tomaszewska et al., 2021b, Tomaszewska et al., 2021a), which likely result in a highly diverse gene pool that can be leveraged for continued improvement through breeding. Exploiting sub-genome variability among species and ploidy levels has been highly successful in the improvement of other crop tribes, such as Triticeae and Brassicaceae (Gale and Miller, 1987, Burton et al., 2004, Ali et al., 2016). However, genetic composition and relationships in *Urochloa* are poorly understood; studies from countries with active *Urochloa* breeding programmes have explored the phylogeny in these species to inform breeding, but projects leveraged of a limited number of markers, such as ITS, RAPD, SSR, ISSR, and microsatellites (Torres González and Morton, 2005, Jungmann et al., 2010, Vigna et al., 2011a, Vigna et al., 2011b, Ferreira et al., 2016, Triviño et al., 2017)

*Urochloa* spp. with apomictic or mixed reproduction have particularly resulted in odd levels of ploidy and contribute to increased intraspecific variability. Polyploidy has many benefits for plants, namely heterosis, gene redundancy, and loss of self-incompatibility and gain of asexual reproduction. In a recent work (Tomaszewska et al., 2021b), we used flow cytometry to determine the ploidy of over 350 *Urochloa* accessions from these collections and propose an evolutionary model. This work extended and corrected some previous studies (Penteado et al., 2000, Mendes-Bonato et al., 2002). We also concluded ploidy was not related to geographical origin, which agrees with previous results (Jungmann et al., 2010, Vigna et al., 2011a, Vigna et al., 2011b, Triviño et al., 2017). In another recent work (Worthington et al., 2021), we have made available a genome assembly and gene annotation of a diploid accession of *U. ruziziensis* (GCA_003016355), which has allowed a greater use of genomics to characterise these materials. For example, we identified loss-of-function (LOF) genes related to forage quality and environmental impact using allele mining (Hanley et al., 2020).

Here, we have characterised the genetic make-up and population structure of 111 accessions, which are representative of the collections of wild materials in Africa in 1984 and 1985. These 111 accessions belong to the five *Urochloa* spp. that are used in the development of commercial forage cultivars. We used RNA-seq from total RNA, so tentatively encompassing the complete *Urochloa* gene pool used in breeding to obtain a definitive picture of the available diversity, resolve questions raised by previous studies, and identify prospective founders to improve the breeding gene pool.

## METHODS

### RNA extraction and sequencing

We sequenced 111 accessions from five *Urochloa* (syn. *Brachiaria*) species. 104 accessions were sampled at the single time from the *in-situ* field collection maintained by the Genebank at the International Center for Tropical Agriculture (CIAT) in Cali, Colombia. Accessions sourced from CIAT are named as e.g. “CIAT 26146”, but we have removed “CIAT” from our text. Fresh leaf material was collected and immediately frozen in liquid nitrogen. Samples were ground in liquid nitrogen and lyophilised. Total RNA was extracted as described in Hanley et al. (2020) with the difference that prior to DNAse treatment the pellets were dried in a rotary evaporator (Eppendorf, USA) and stored at room temperature. Another seven accessions were obtained from the United States Department of Agriculture (USDA, GA, USA) as seeds. These seven accessions include “PI” at the beginning of their ID. These seven accessions were sampled at a different time than the other accessions after growing in glasshouses at University of Leicester, UK. We generated one single sample from each accession, and we use “sample” and “accessions” as synonyms in our case through the text. For all samples, Illumina sequencing using standard RNA-seq library preparations with 150 bp paired reads was conducted by Novogene Europe (Cambridge, UK). The raw reads were deposited in SRA under Bioproject PRJNA513453.

### Read alignment and SNP calling

Raw reads were pre-processed using Trim galore v. 0.5 (Krueger, 2015) with the options for Illumina paired reads and trimming 13 bps at the 5’ end in both reads. Processed reads were aligned to the available *Urochloa* genome (Worthington et al., 2021), which corresponded to a *U. ruziziensis* accession. RNA to DNA alignments were done using STAR v. 2.6.0c (Dobin et al., 2013) with a minimum overlap of 30 % and a maximum mismatch of 3 bp per alignment, in order to allow for mapping from more distant species to the genome. Alignment coverage was calculated using BEDTools genomecov. SNP calling was done using GATK v. 3.7.0 and the recommended pipeline for RNA-seq (Van der Auwera et al., 2013). Firstly, we used PicardTools v. 2.1.1 to annotated duplicate reads using the option MarkDuplicates. Later, we used GATK’s tool SplitNCigarReads with the options “-rf ReassignOneMappingQuality -RMQF 255 -RMQT 60 -U ALLOW_N_CIGAR_READS” to reformat some alignments that span introns to match conventions for the final step. The final step was SNP calling using GATK’s tool HaplotypeCaller with all the samples at the same time (multisample mode). SNP calling was run with the options “-ploidy 6 -dontUseSoftClippedBases -stand_call_conf 20 - maxNumHaplotypesInPopulation 128” to obtain a good quality calling from RNA alignments. GATK identified 6,461,493 variants, which included 5,757,116 SNPs. These were filtered for a minimal allele frequency (MAF) of 1% to give a set of 4,722,195 SNPs. Sites with a depth of lower than 5 were set to missing, then sites with more than 40% missing data were removed to give a final set of 1,167,542 SNPs. Two additional subsets were obtained by filtering out either the 67 samples (895,667 SNPs) in the agamic group or the *U. humidicola* samples (512,611 SNPs). These subsets were filtered for MAF of 1%.

### Population analysis

Population structure analysis was performed through ADMIXTURE (Alexander and Lange, 2011) using *K* = 3 to *K* = 10 for the 111 samples, *K* = 2 to *K* = 8 for the 67 samples and *K* = 2 to *K* = 8 for the 28 samples. Each value of *K* was run 10 times, the cross-validation error was averaged over the 10 runs. The 10 output files were combined using CLUMPP within the R package POPHELPER v.2.2.7 (Francis, 2017). The PCA (principal component analysis) was carried out using Tassel v5.2.41 (Bradbury et al., 2007).

## RESULTS

### Sequencing, aligning and SNP calling in a panel of *Urochloa* accessions from five species

We sequenced 111 accessions from five *Urochloa* (syn. *Brachiaria*) species: *U. ruziziensis*, *U. brizantha*, *U. decumbens, U. humidicola* and *U. maxima* (syn. *Megathyrsus maximus*). Species identity and ploidy were previously determined using plant architecture traits and flow cytometry of fluorescently stained nuclei (Tomaszewska et al., 2021a, Tomaszewska et al., 2021b). The country of origin of 92 accessions was known and for 75 accessions we also knew the collection coordinates (Fig. 1). Accessions were collected in a broad range of latitudes (20.08S to 11.37N) but not of longitudes (26.98E to 42.05E), except for one *U. brizantha* accession from Cameroon. Annotations were summarised in Table 1 and detailed in Suppl. Table 1.

**Figure 1.**
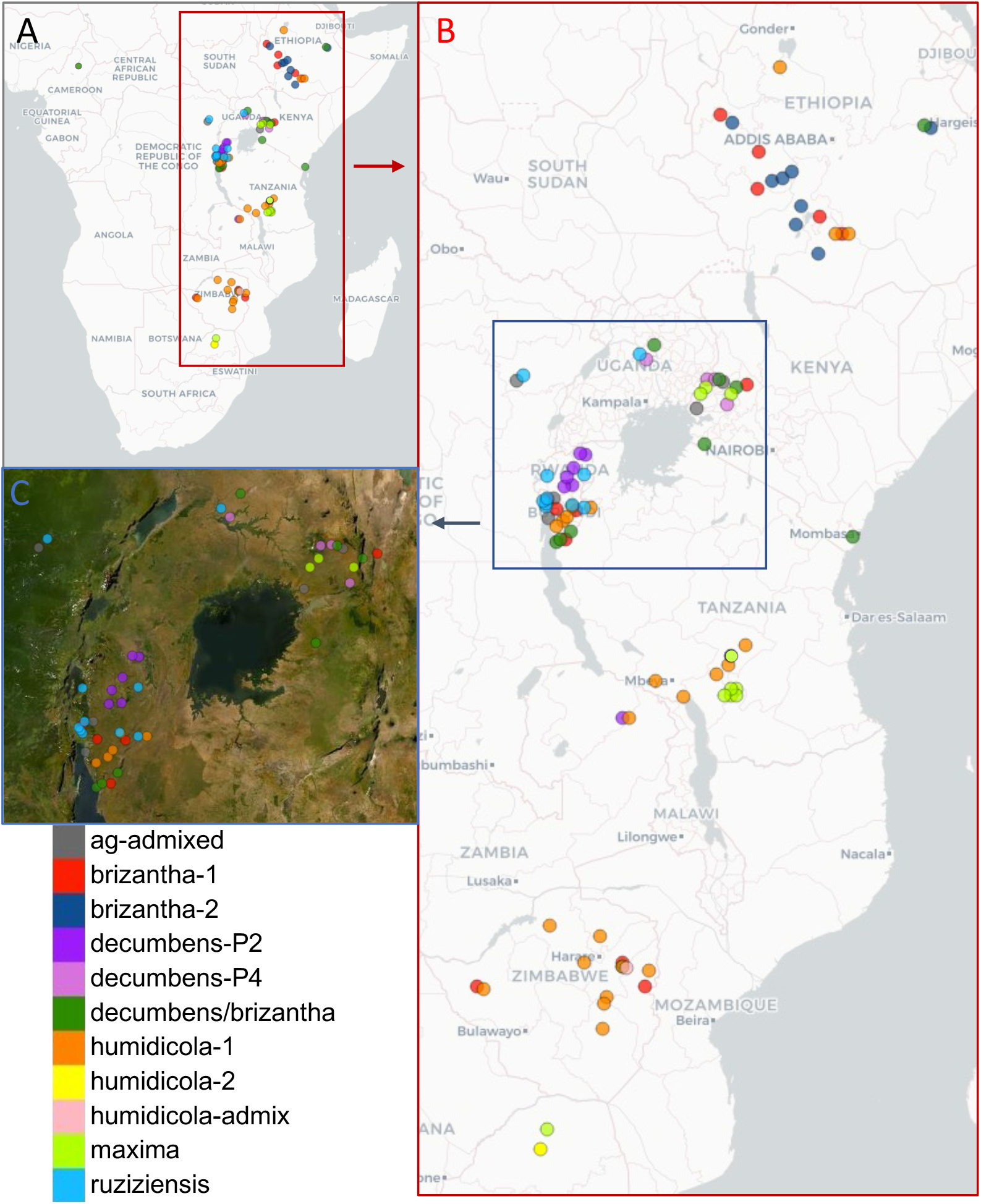
Geographical origin of 92 accessions with collection coordinates (74 accessions) or country of origin (18 accessions). Accessions were coloured by subpopulation. (A) Origin in Sub-Sahara Africa. (B) Zoom into East Africa. (C) Zoom into the Great Lakes region.

**Table 1:** Summary of the species (rows) and ploidy level (columns) of the 111 accessions used in this study. Details about each accession were included in Suppl. Table 1.

|  | 2n | 4n | 5n | 6n | 7n | 8n/9n | ? | Total |
| --- | --- | --- | --- | --- | --- | --- | --- | --- |
| *U. ruziziensis* | 10 |  |  |  |  |  | 1 | 11 |
| *U. decumbens* | 8 | 17 |  |  |  |  | 1 | 26 |
| *U. brizantha* |  | 17 | 9 | 2 |  |  | 1 | 29 |
| *U. humidicola* |  |  |  | 10 | 16 | 2 |  | 28 |
| *U. maxima* |  | 12 |  |  |  |  | 1* | 13 |
| *U. spp.* |  | 2 | 1 |  |  |  | 1 | 4** |

\*Accession 26438 was received as *U. humidicola* but it was reassigned to *U. maxima* based on the results. \*\*Accession 1752 (*U. spp.*) clustered within the agamic group.

Samples were aligned to the available *Urochloa* genome assembly and annotation (Worthington et al., 2021), which corresponds to a diploid *U. ruziziensis* sample. Two well-defined groups of species were observed based on aligning metrics (Fig. 2); over 70 % of the reads from *U. ruziziensis*, *U. decumbens* and *U. brizantha* (all but one) accessions had more than 70 % of reads that aligned in the reference genome once (uniquely-mapping reads). On the contrary, accessions from *U. maxima* and *U. humidicola* showed a percentage of uniquely-mapping reads under 70 % (Fig. 2A). The grouping was correlated to the genetic distance to the reference genome (reference bias).

**Figure 2.**
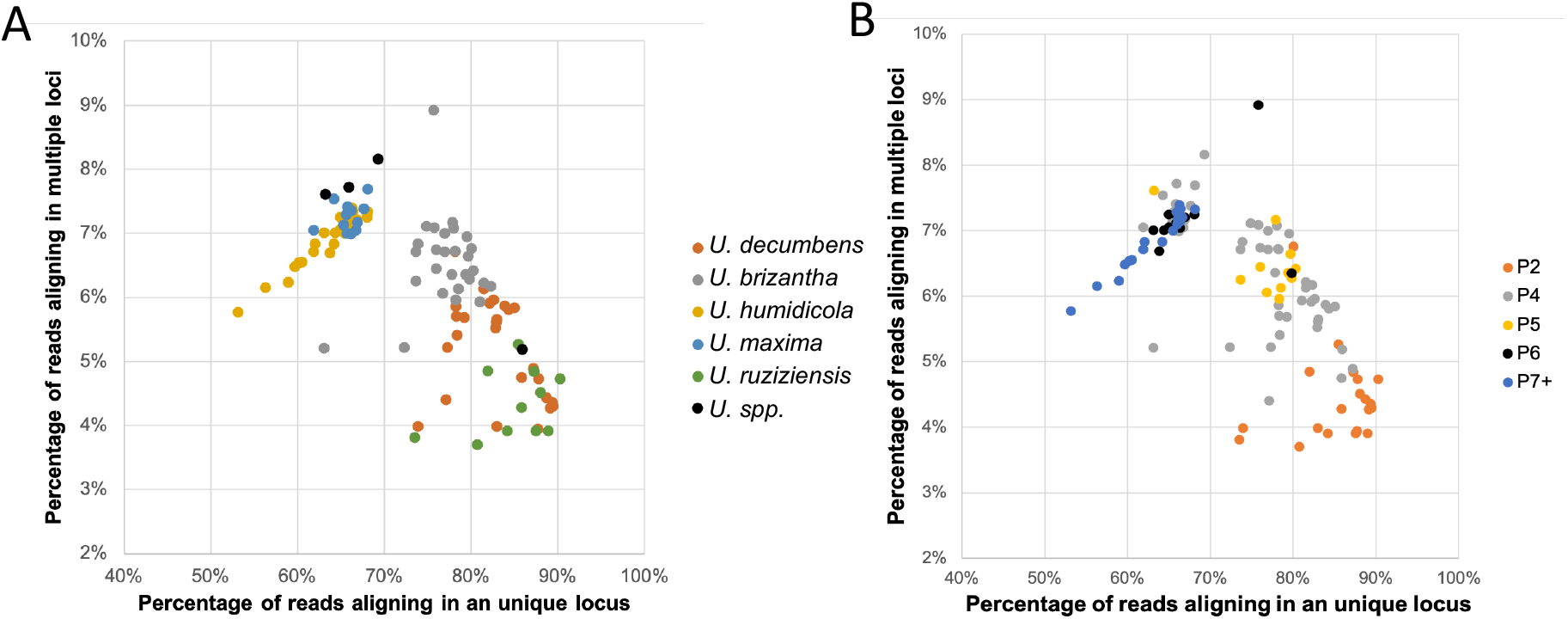
Percentage of reads aligning in either uniquely or in multiple positions in the genome. The 111 accessions were coloured by species (A) or ploidy (B).

The percentage of reads mapping in multiple loci (multi-mapping reads) increased with ploidy (Fig. 2B) for the group of the accessions belonging to the species *U. ruziziensis*, *U. decumbens* and *U. brizantha*; diploids had a percentage of multi-mapping reads under 5 %, while it was over 5 % in most polyploid accessions. However, the percentage of multi-mapping reads in the other species, which are more distant species to the reference, was directly proportional to the total number of mapped reads (Fig. 2B), i.e. independent of ploidy.

RNA-seq reads covered 268.84 Mbps (~36.7 % of the 732.5 Mbps genome assembly). The covered regions are more than 2.5 times the original gene annotation from the *U. ruziziensis* genome (43,152 genes comprising 102 Mbps). The median read coverage was 25 reads in the covered regions, and the average read coverage in these regions was 2587 ± 54293 reads. After SNP calling and filtering, the average SNP density in the genome was 7.3 SNPs/Kbp, or 17.9 SNPs/Kbp if only considering the read covered part of the genome. Using the 43,152 genes and 202,681 exons annotated in the genome reference, the median was 69 and 13 SNPs in each gene and exon (average was 95 and 36 per gene and exon, respectively). 34,981 of the annotated genes had at least one SNP.

### Admixture analysis

We employed genetic admixture analysis for defining subpopulations. To assign 111 *Urochloa* accessions to subpopulations, the admixture (Fig. 3) and principal component (Fig. 4) analysis were considered together. The “admixture model” assumes that each individual has ancestry from one or more of “K” genetically distinct sources. An estimation of four subpopulations (K = 4) was selected based on the CV error (Suppl. Fig. 1A) and population structure (Fig. 4). A minimum threshold of 50% genetic composition was used to assign accessions to groups. This allowed us to place the accessions in four groups (Fig. 3): *U. humidicola* (28 accessions), *U. maxima* (13 accessions), “agamic group 1” (54 accessions from the three remaining species) and a closely related “agamic group 2” (that corresponded with the “brizantha-1” subpopulation). Three samples obtained from USDA and identified simply as “*Urochloa* sp.” showed an admixture of these four groups and were annotated as “admixed”. Accession 26438 (sample 86) was received as *U. humidicola*. Since it clustered with the *U. maxima* accessions, we reassigned it into that species. When we reduced the number of groups (K = 3), the *U. humidicola* and *U. maxima* species clustered together, but the agamic groups 1 and 2 were consistent (Suppl. Figure 2). When we increased the number of groups (K = 5), a new group split from the “agamic group 1” (that corresponded with the “brizantha-2” subpopulation). The twenty-eight accessions in the *U. humidicola* group had a basic chromosome number of 9 and high ploidy levels ranging from 6 to 9. The twelve accessions in the *U. maxima* group had a basic chromosome number of 8 and are tetraploid. The 67 accessions in the agamic groups had a basic chromosome number of 9 and ploidy levels ranging from 2 to 6 (Tomaszewska et al., 2021b).

**Figure 3.**
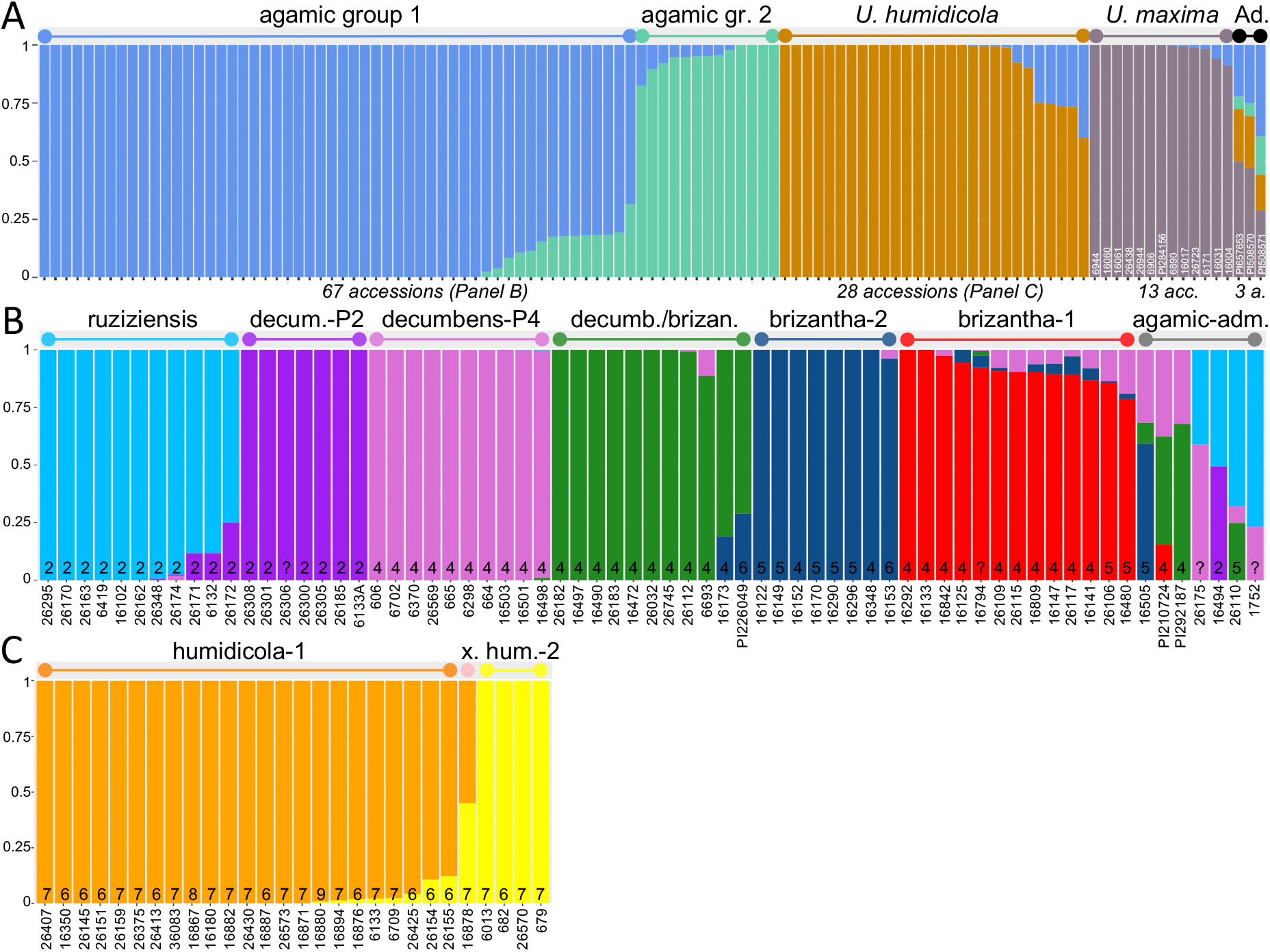
Admixture analysis of the genetic ancestry inferred in the complete set of 111 accessions (A), the subset of 67 accessions in the agamic group (B), and the subset of 28 *U. humidicola* accessions (C). Ploidy level is included at the foot of each column. Each accession is represented by a stack column partitioned by proportion of ancestral genetic component, where each identified ancestral genetic component is represented with a different colour. Accessions with a single colour are “pure”. A minimum threshold of 50 % (A) or 70 % (B and C) genetic composition was used to assign accessions to groups. In panel C, “x” for “humidicola-admix”.

**Figure 4.**
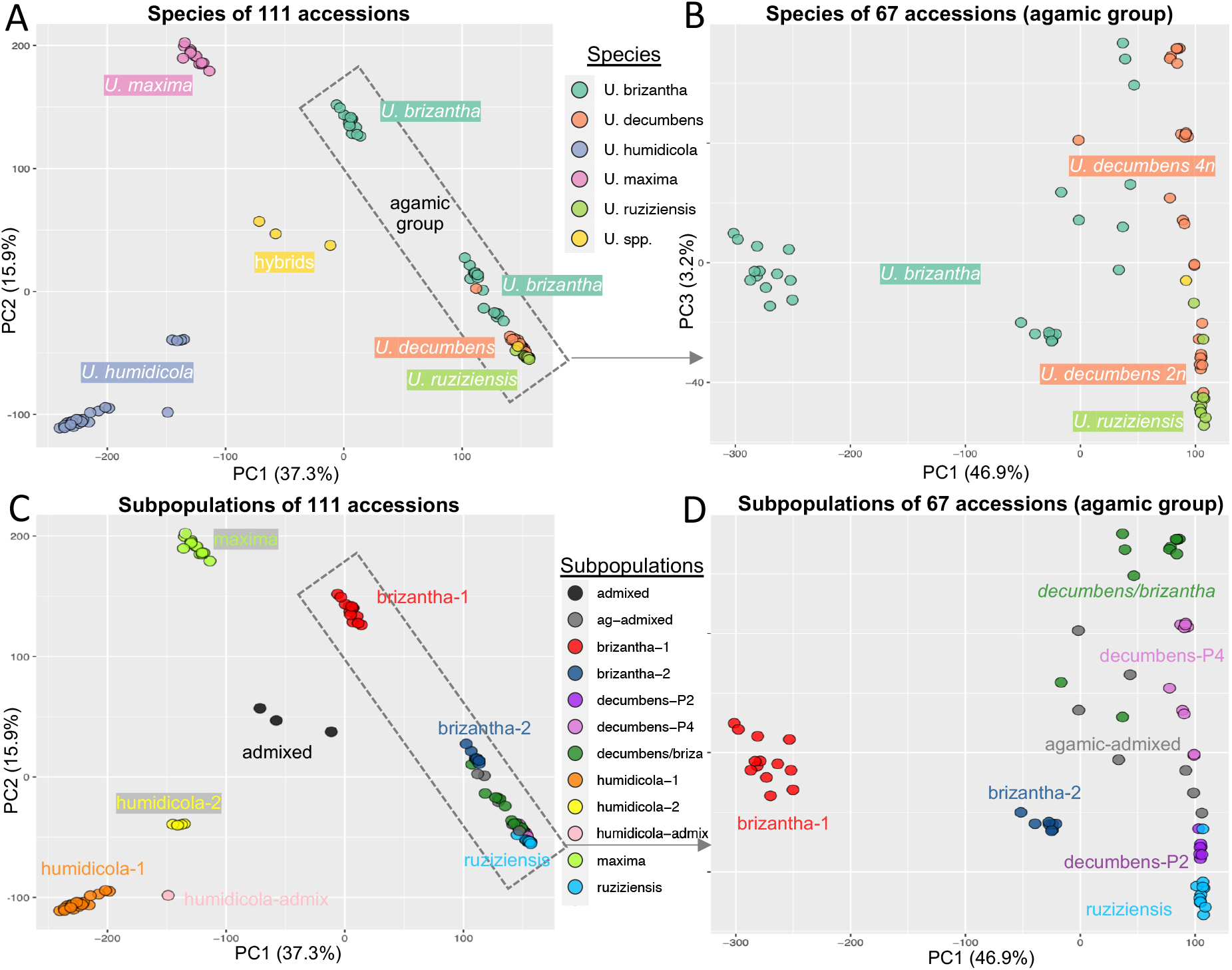
Population structure by Principal Component Analysis (PCA) using the top two components to separate the complete set of 111 accessions (A and C) or the components 1 and 3 to separate the subset of 67 accessions in the agamic group (B and D). Accessions were coloured by species (A and B) or subpopulation (C and D).

The admixture analysis was subsequently carried out using only the 67 accessions in the agamic group (Fig. 3B). An estimation of six groups (K = 6) was selected based on the CV error (Suppl. Fig. 1B) and population structure (Fig. 4). A minimum threshold of 70 % shared genetic composition was used to assign accessions to each of the six groups. As shown in fig. 3B, the group “ruziziensis” was composed of all eleven *U. ruziziensis* accessions. It included one accession wrongly classified as *U. decumbens*. Within it, five samples showed shared ancestry (1-25%) with diploid *U. decumbens*. All seven diploid *U. decumbens* accessions composed the group “decumbens-P2” and were pure accessions with no shared ancestry with any other group. Similarly, ten tetraploid *U. decumbens* formed the group “decumbens-P4” with pure accessions with no shared ancestry with any other group. However, another six tetraploid *U. decumbens* composed a different group together with five *U. brizantha* accessions, which was called “decumbens/brizantha”. This group of eleven accessions was the only one composed by more than one species. Despite this mix, these accessions showed clear shared ancestry among them and no shared ancestry with any other group (except two samples with minor components). Finally, the group “brizantha-1” and “brizantha-2” were formed by eight and thirteen *U. brizantha* accessions, respectively. The group “brizantha-2” has pure accessions with no shared ancestry with other groups (with one minor exception under 5%); while most samples in “brizantha-1” have shared ancestry with “decumbens-P4”. The group “brizantha-1” corresponds to the previous “agamic group 2”. The “brizantha-2” subpopulation was only observed in Ethiopia, while “brizantha-1” was observed in a broad range of latitudes. When we reduced the number of groups (K = 5), the “brizantha-decumbens” merged with the “decumbens-P4”. When we increased the number of groups (K = 7), five “brizantha-1” split into an independent subpopulation (Suppl. Fig. 3).

The admixture analysis was finally completed using only the twenty-eight *U. humidicola* accessions (Fig. 3C). An estimation of two groups (K = 2) was selected based on the CV error (Suppl. Fig. 1C) and population structure (Fig. 4). A minimum threshold of 70 % shared genetic composition was used to assign accessions to a group. The twenty-eight samples into the two groups: 23 accessions into “humidicola-1” and four accessions into “humidicola-2”. Accession 16878 was an equal mixed from both *U. humidicola* groups and annotated as “humidicola-admixed”. When we increased the number of groups (K = 3 and K = 4), we obtained a small subpopulation with the accessions with higher admixture (16878 and 26155) and an artificial split with some “humidicola-1” accessions in an additional group (Suppl. Fig. 4).

A smaller number of thirteen *U. maxima* accessions showed little genetic diversity compared to the other species. Because of the low diversity, we assigned all the *U. maxima* to a single subpopulation, named “maxima”.

### Population structure by principal component analysis

A principal component analysis (PCA) showed the relationship between the 111 accessions, species and admixture groups (Fig. 4A and 4C). The PCA was also done for the 67 accessions in the agamic group alone (Fig. 4B and 4D). The PCA analysis allowed us to define 12 clusters in total, which easily corresponded with the 10 subpopulations and two admixed groups. The distribution of accessions into subpopulations according to the species and ploidy annotations is represented in figure 5.

**Figure 5:**
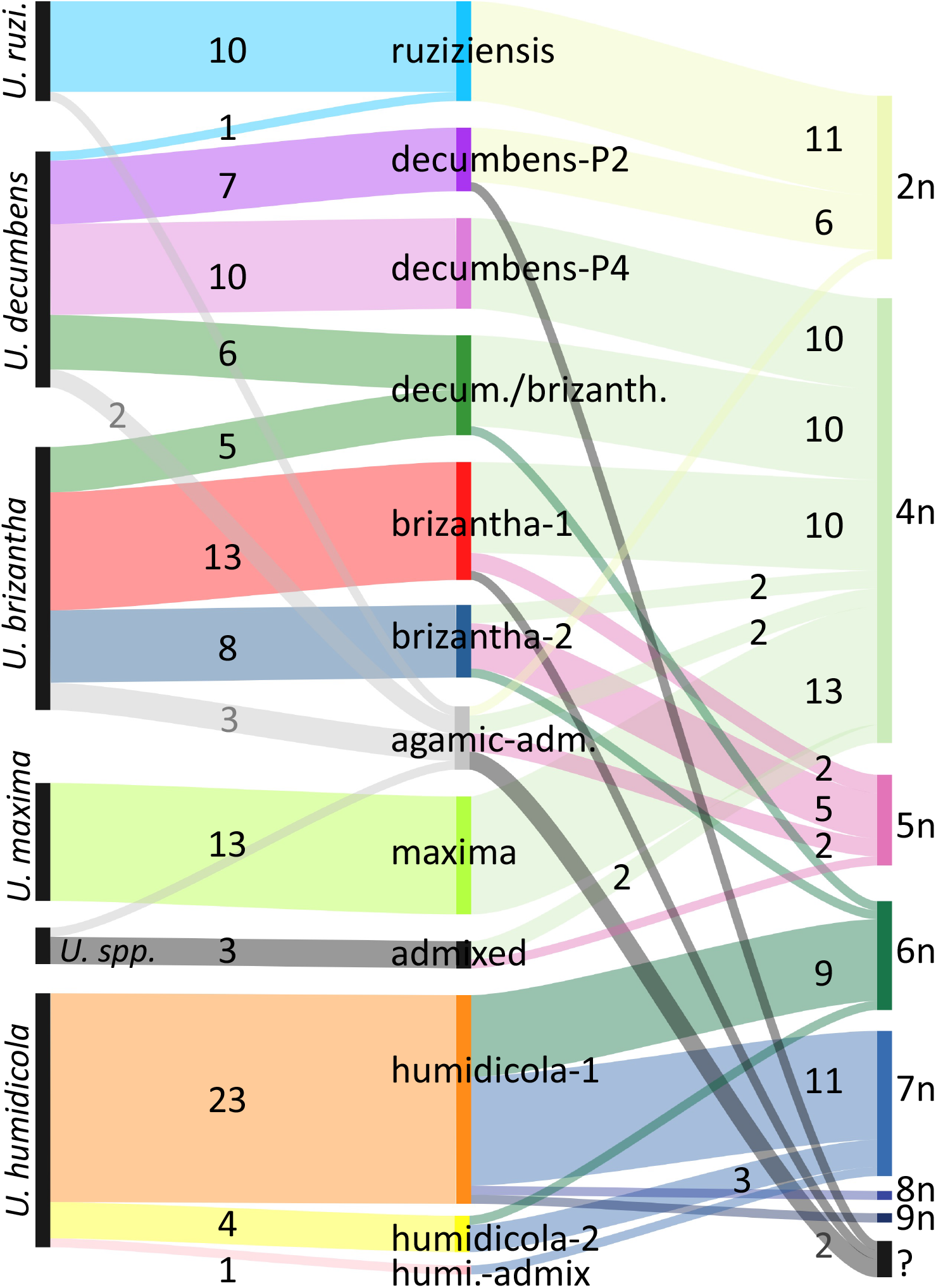
Distribution of accessions in subpopulations according to the species and ploidy annotations. The number on each stream represents the number of accessions in that division. Streams without a number represent a single accession.

All subpopulations contained accessions from a single species, except subpopulation “decumbens/brizantha”. Remarkably, this subpopulation contained samples that showed greater similarity to each other -in spite of species- than to accessions from the same species in different subpopulations. The two diploid subpopulations, “decumbens-P2” and “ruziziensis” clustered together and apart from polyploid subpopulations. Subpopulation “brizantha-1” was more distant to other agamic subpopulations than “brizantha-2” despite accessions in “brizantha-1” showed shared admixture with tetraploid *U. decumbens*, while accessions in “brizantha-2” did not.

Two groups of accessions contained hybrids, one containing hybrids between the distant *Urochloa* species (“admixed” subpopulation) and another contained hybrids within the three species in the agamic group (“agamic-admixed”), which readily interbreed in control conditions.

## DISCUSSION

We clarified the relationship between the gene pools from five *Urochloa* spp. that are used in the development of commercial forage cultivars. By using RNA-seq, we leveraged in an unprecedented number of markers, over 1.1M SNPs, that virtually encompassed the complete transcriptome from the accessions based on the total genome length covered by the reads (~ 269 Mbp or 37 % of the genome). We obtained a median of 69 and 13 SNP sites per gene and exon, respectively, which makes this dataset a valuable resource for breeders and researchers (e.g. to design screening markers). The greater length compared to the annotated gene models (Worthington et al., 2021) was probably because the later only used transcriptomic data from *U. ruziziensis*. However compared with our dataset, that transcriptomic data was from multiple tissues and included stress conditions. The genus *Urochloa* includes species previously classified under other taxonomic groups. We have opted to annotate all as *Urochloa* supported by recent work (Tomaszewska et al., 2021b). E.g., we did observed the same distance between *U. maxima* and the agamic group than between *U. humidicola* and the agamic group.

We observed two subpopulations of *U. brizantha*, two subpopulations of *U. decumbens*, and one subpopulation with accessions from both *U. brizantha* and *U. decumbens*. The two *U. decumbens* subpopulations were divided by ploidy. Diploid *U. decumbens* clustered with *U. ruziziensis*, while tetraploid *U. decumbens* clustered with *U. brizantha*. This split in two *U. decumbens* subpopulations by ploidy was previously reported using microsatellites (Triviño et al., 2017). In previous studies, the relation of *U. decumbens* with the other two species has been discussed, as it was alternatively found closely related to *U. ruziensis* (Ferreira et al., 2016) or *U. brizantha* (Ambiel et al., 2008). In fact, both observations were correct depending on the ploidy of the accessions under consideration.

Based on our results, *U. brizantha* diversity is complex and divided in several gene pools. A group of eleven *U. brizantha* accessions was different enough to the rest of the agamic group to form an independent cluster (“agamic group 2”; Fig. 3A and 4A). This group of eleven accessions later formed the subpopulation “brizantha-1”. Despite of “brizantha-1” being distant, we observed admixture between “brizantha-1” and “decumbens-P4”, “brizantha-2” and “decumbens/brizantha” subpopulations (Fig. 3B). Among the possible evolutive scenarios that would explain the multiple shared ancestry in “brizantha-1” despite being more distant, previous studies have proposed a single polyploidization event taking place to establish both the tetraploid *U. brizantha* and *U. decumbens* (Pessoa-Filho et al., 2017, Tomaszewska et al., 2021b). The “brizantha-1” subpopulation was observed in a broad range of latitudes (e.g. in Ethiopia and Zimbabwe), while “brizantha-2” was only observed in Ethiopia.

We obtained a subpopulation, named “decumbens/brizantha”, that included an almost equal number of *U. decumbens* and *U. brizantha* accessions. This is the only subpopulation with more than one species, and most accessions did not show shared ancestry with the other subpopulations from either of these species (Fig. 3B). Furthermore, the PCA also showed “decumbens/brizantha” clustered independently to other groups (in the top right corner of Fig. 4D). Remarkably, the “decumbens/brizantha” merged with the “decumbens-P4” when we did the admixture analysis with less subpopulations (K = 5). At the same time, two accessions (16173 and PI226049) shared ancestry with “brizantha-2” and were situated between the subpopulations “decumbens/brizantha” and “brizantha-2” in the PCA (Fig. 4D). Consequently, we concluded “decumbens/brizantha” cannot be merged with either “decumbens-P4” or “brizantha-2”, but on the contrary, evidence supported it is an independent subpopulation.

Vigna et al. (2011b) observed three clusters in *U. brizantha* after evaluating 172 accessions from EMBRAPA’s collection (so resulting from the same field work in 1980s than our dataset) using 20 SSR markers. Eleven accessions are common between both studies, and our subpopulations “brizantha-1” and “brizantha-2” corresponded with clusters II and I, respectively (Vigna et al., 2011b). Notably, their cluster III appears to include additional “brizantha-1” and “brizantha-2” accessions (16122, 16480), so does not correspond with our “decumbens-brizantha” subpopulation. Triviño et al. (2017) did not discussed a division among *U. brizantha* accessions, but included a tree resulting from UPGMA clustering based on 39 microsatellites that would also support at least two gene pools in *U. brizantha*.

In the centre of the agamic group, we identified the “agamic-admixed” accessions (Fig. 4D). This cluster of accessions included hybrid accessions resulting from interspecific species within the agamic group, and should not be confused with the “admixed” accessions (Fig. 4C), which resulted from crosses between more distant *Urochloa* species. Our analysis supports that the “agamic-admixed” are either progeny from crosses between *U. decumbens* and either *U. decumbens* or *U. brizantha* (16505, PI210724, PI292187); or between *U. ruziziensis* and either *U. decumbens* or *U. brizantha* (26175, 16494, 26110, 1752).

*U. maxima* is also known as *Panicum maximum* or *Megathyrsus maximus*. All *U. maxima* accessions (including accession 26438, which was incorrectly annotated as *U. humidicola*) showed very limited diversity (Fig. 4) and were assigned to a single subpopulation (“maxima”). This could reflect lower diversity in the species, or be a consequence of original collection and sampling strategy, but it suggests there would be limited gains from including multiple accessions from our study in breeding programmes.

We observed two different subpopulations in *U. humidicola*, named “humidicola-1” and “humidicola-2”, plus a single accession (16878) that was an equal mix from both subpopulations. In the initial admixture analysis with all the 111 accessions (Fig. 3A), the “humidicola-2” and “humidicola-admixed” accessions had shared ancestry to the “agamic group 1”, while the “humidicola-1” did not. Triviño et al. (2017) also observed a large group of *U. humidicola* accessions including all but three of their accessions. These three distant *U. humidicola* accessions were 675, 679 and 26146. Accession 679 is a “humidicola-2” subpopulation in our study. Remarkably, 26146 is the sexual *U. humidicola* accession that allowed the establishment of breeding programmes in mid-2000s, and combining the results from both studies supports that the sexual 26146 accession is likely a “humidicola-2” accession. We only had collection information for one “humidicola-2” accession, but it was close to a “humidicola-1” accession, so it is not likely a geographical division. Vigna et al. (2011a) analysed 26 *U. humidicola* accessions and used UPGMA clustering based on 38 microsatellites to divide *U. humidicola* in two branches in the resulting tree. All seven common accessions with our study were “humidicola-1” and appeared in the top branch of the tree. The bottom branch may be “humidicola-2”, since it included the sexual accession 26146, one accession (26149) not sequenced in our dataset, and the progeny from their crossing. On the other hand, two studies from the same group (Jungmann et al., 2010, Vigna et al., 2011a) identified five clusters in *U. humidicola* accessions using ~ 50 SSR markers. While multiple accessions were common to both studies, we did not find correlation between our results and the five clusters; i.e. “humidicola-1” accessions were evenly divided among the multiple clusters. In a similar conclusion, when we increased the number of tentative *U. humidicola* subpopulations (K = 3 and K = 4), we found the additional groups to be artificial splits from “humidicola-1” and not supported by the PCA.

## CONCLUSION

We clarified the relationship between the gene pools from five *Urochloa* spp. that are used in the development of commercial forage cultivars in different countries. We identified ten subpopulations in total, which had no relation with geographical collection, and represent ten independent gene pools (excluding the two admixed subpopulations). Our results support the division in *U. decumbens* by ploidy, with a diploid subpopulation closely related to *U. ruziziensis*, and a tetraploid subpopulation closely related to *U. brizantha*. We observed clearly differentiated gene pools in *U. brizantha*, which were not related with origin or ploidy. One of these gene pools, named “brizantha-1”, clustered relatively distant to the rest of the agamic accessions despite having significant shared ancestry with tetraploid *U. decumbens*. Among the possible evolutive scenarios to explain this observation, it would support a single polyploidization event taking place to establish both the tetraploid *U. brizantha* and *U. decumbens*. The “brizantha-1” gene pool should be further explored for prospective founders in the agamic group since *U. brizantha* is particularly tolerant to neotropic insects, which is one of the main traits under selection. We also identified a well-supported subpopulation containing both polyploid *U. decumbens* and *U. brizantha* accessions that likely constitutes a third independent gene pool for both species. We observed two gene pools in *U. humidicola*. One subpopulation, “humidicola-2”, was significantly smaller but likely includes the only known sexual accession. We also observed one case of natural hybridization between both *U. humidicola* groups. Our results offer a definitive picture of the available diversity and resolve questions raised by previous studies. They provide an insight into the diversity available for improvement through crossing, and a platform to identify target genes for forage grass improvement, also providing gene sequences to allow for genome editing (CRISPR/Cas9) approaches. Furthermore, as performance data become available, the data could be further leveraged for GWAS (Genome Wide Association Studies), genotyping array construction, and development of genetic markers for selection in breeding programmes.

## AUTHOR CONTRIBUTIONS

JJDV, RACM, JT, TS and JSHH conceived and managed the project. JH and JJDV completed the bioinformatics analysis. PT analysed and provided ploidy level and accessions information. TKP, VC and JA selected, validated and collected the samples. TKP carried out RNA extraction. JH and JJDV wrote the manuscript with contributions from all the authors.

## ACKNOWLEDGEMENTS

This work was supported under the RCUK-CIAT Newton-Caldas Initiative “Exploiting biodiversity in *Brachiaria* and *Panicum* tropical forage grasses using genetics to improve livelihoods and sustainability”, with funding from UK’s Official Development Assistance Newton Fund awarded by UK Biotechnology and Biological Sciences Research Council (BB/R022828/1). Additional funding for this study was received from the CGIAR Research Programmes on Livestock; and Climate Change, Agriculture and Food Security (CCAFS). JH and JJDV received additional funding from the Biotechnology and Biology Sciences Research Council (BBSRC)’s Global Challenge Research Fund (Project BB/P028098/1) and core strategic funding (Project BBS/E/T/000PR9818).

We are grateful to CIAT’s Genebank and USDA’s Germplasm Resources Information Network (GRIN) for their generous provision of germplasm. Germplasm held in the CIAT and USDA collections is available on request.

## DATA AVAILABILITY

All the raw reads were deposited in SRA under Bioproject PRJNA513453.

---

Supplementary Table 1. Sample number, accession number, species, ploidy, subpopulation, architecture, collection location, and PCA position for each of the 111 accessions used in this study.

Supplementary figure 1: Cross-validation (CV) error and chosen value for number of groups (K) for the complete dataset of 111 accessions (A), the subset of 67 accessions in the agamic group (B), and the subset of 28 *U. humidicola* accessions (C). Cross-validation error is shown on the Y-axis (vertical) and the number of hypothetical populations on the X-axis (horizontal).

Supplementary figure 2: Admixture analysis for alternative values for number of groups (K = 3, 4 -selected-, and 5) in the complete set of 111 accessions. Numbered by “sample id”.

Supplementary figure 3: Admixture analysis for alternative values for number of groups (K = 5, 6 -selected-, and 7) in the subset of 67 accessions in the agamic group. Numbered by “sample id”.

Supplementary figure 4: Admixture analysis for alternative values for number of groups (K = 2 -selected, 3 and 4) in the subset of 28 *U. humidicola* accessions. Numbered by “sample id”.

