## Supplementary figures and images for "The five *Urochloa* spp. used in development of tropical forage cultivars originate from defined subpopulations with differentiated gene pools"

### Supplementary Figure 1

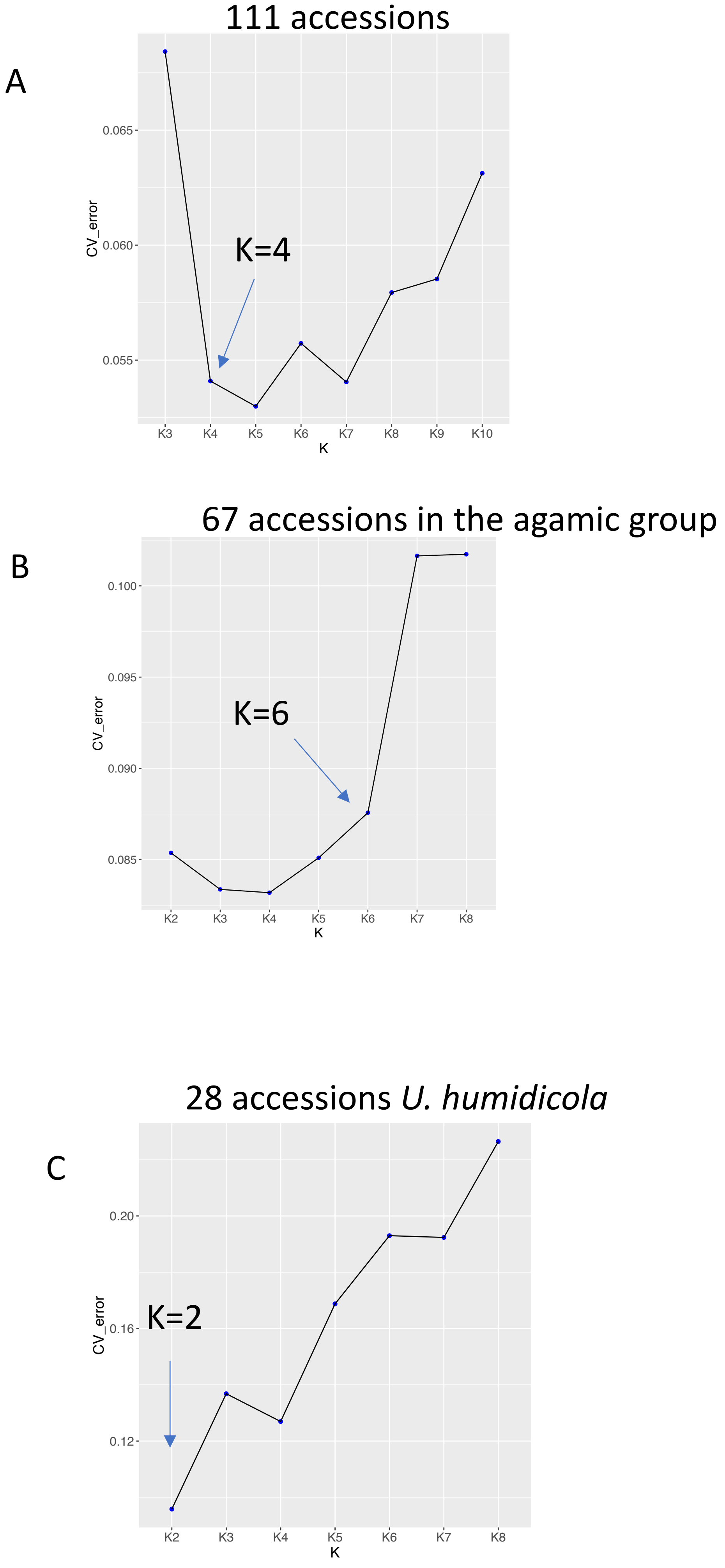

### Supplementary figure 2

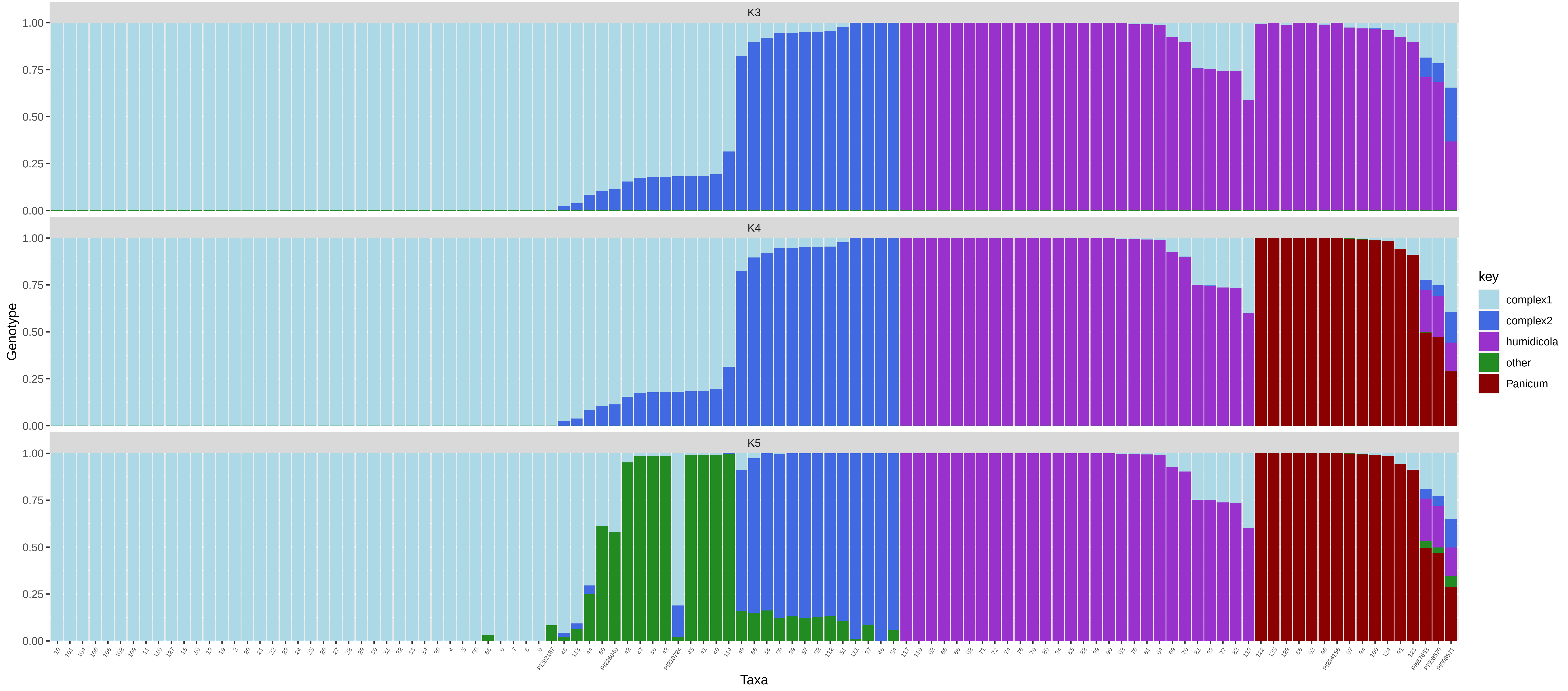

### Supplementary figure 3

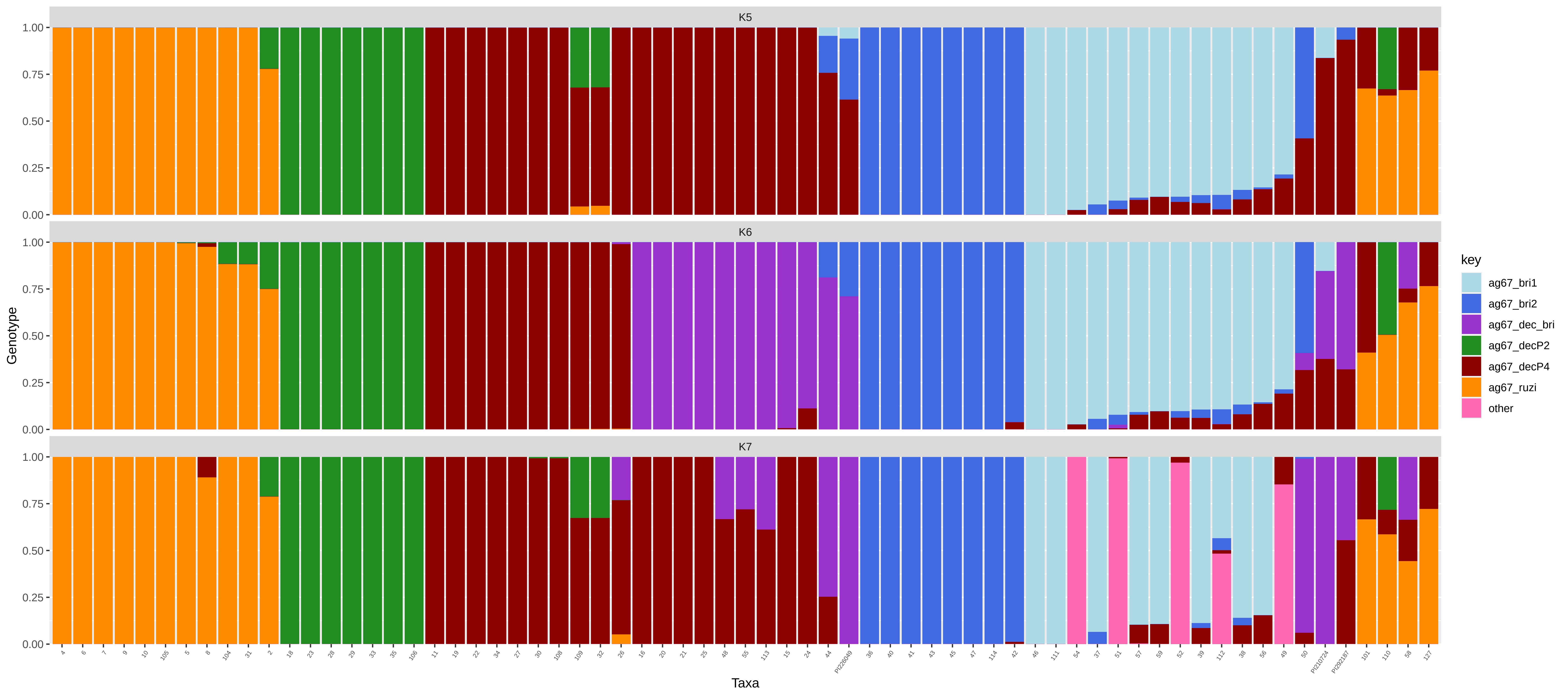

### Supplementary figure 4

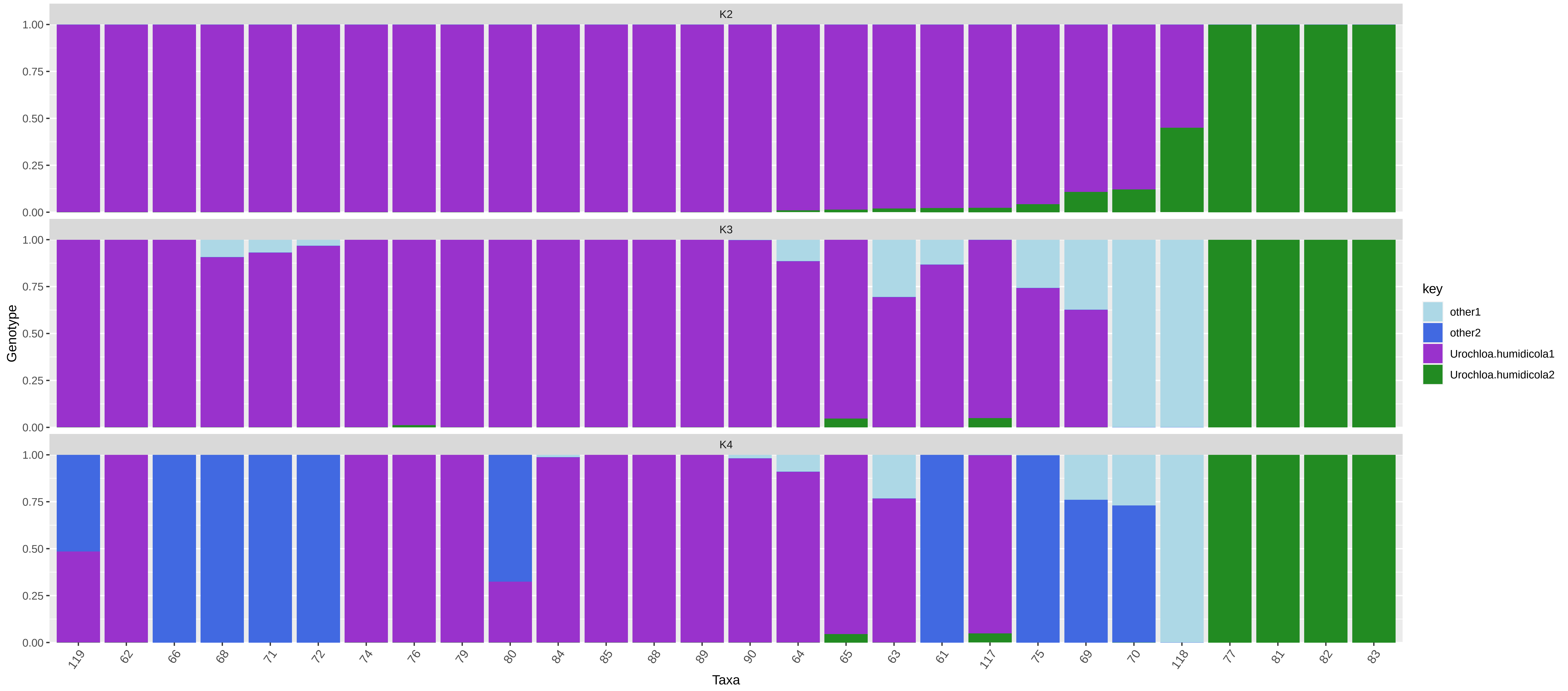
